## Supplementary Figures for "Knowledge-graph-based cell-cell communication inference for spatially resolved transcriptomic data with SpaTalk"

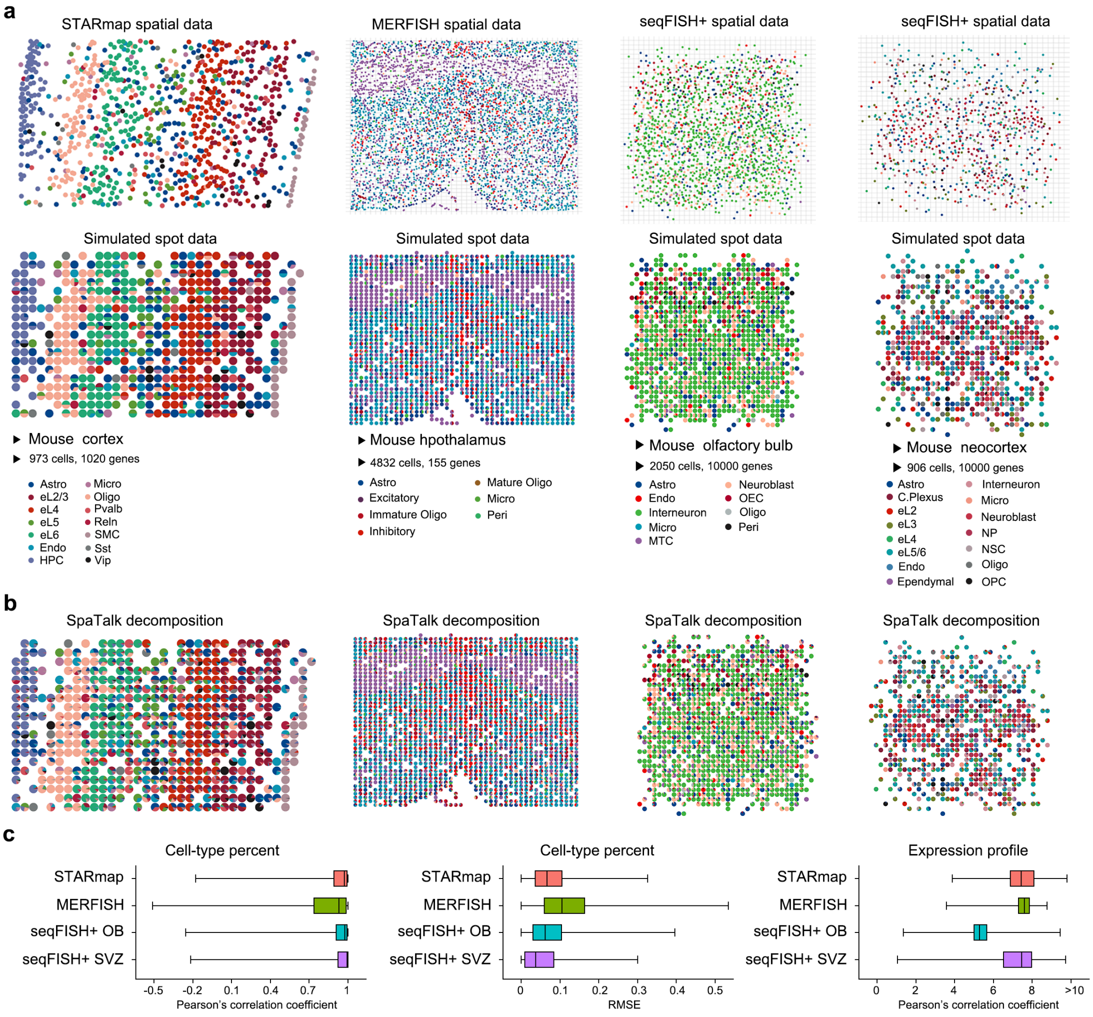


**Fig. 1 Benchmarked datasets and cell-type decomposition by SpaTalk. a** Selected three spatial technologies and four ST datasets at single-cell resolution, namely the STARmap mouse cortex, MERFISH mouse hypothalamus, seqFISH+ OB and SVZ datasets. **b** Cell-type decomposition by SpaTalk over the benchmarked datasets. **c** Performance of SpaTalk over the benchmarked datasets in cell-type decomposition. Pearson's correlation coefficient and RMSE were used to evaluate the predicted and real cell-type percent as well as the expression profile between the spot and the predicted optimal cellular combination for each simulated spot.

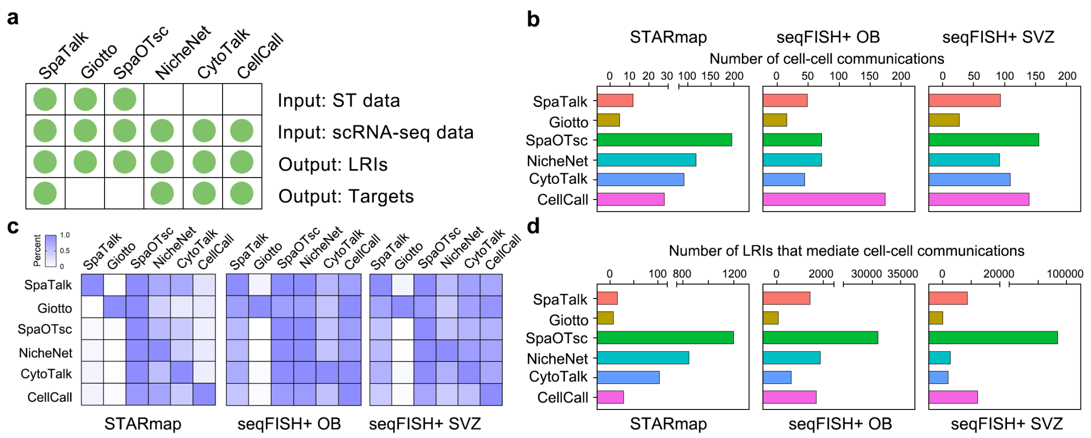


**Fig. 2 Comparison of SpaTalk with other methods. a** Selected representative cell-cell communication inference method, namely the Giotto, SpaOTsc, NicheNet, CytoTalk, and CellCall. Requirements of input data and the type of output data were shown. **b** umber of cell-cell communications inferred by different methods over the STARmap, seqFISH+ OB and SVZ ST data at single-cell resolution. **c** Percent of shared cell-cell communications between paired methods. For each row, the value means the number of overlapped cell-cell communication divided by the number of inferred cell-cell communications for this method. **d** umber of LRIs that mediate cell-cell communications inferred by benchmarked methods over the STARmap, seqFISH+ OB and SVZ ST datasets.

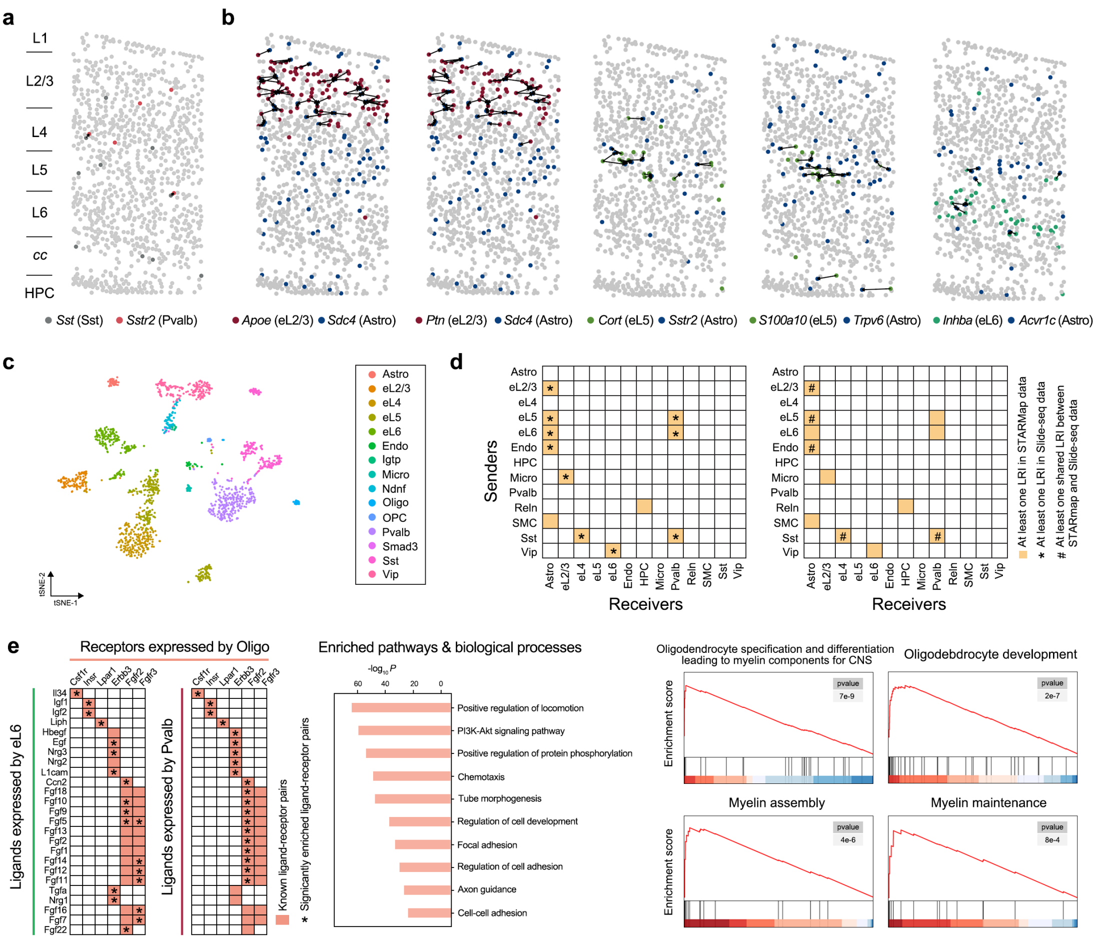


**Fig. 3 Cell-cell communication inference on the STARmap and Slide-seq datasets by SpaTalk. a** direct cell-cell communication mediated by the Sst-Sstr2 interaction between Sst neurons and Pvalb neurons. **b** Inferred LRIs that mediate cell-cell communications among Astro, eL2/3, eL5, and eL6. **c** A previously published adult mouse cortical cell taxonomy by scRNA-seq data, which contains 1718 single cells involving 15 cell types. Astro, astrocytes; eL2/3, eL4, eL5, eL6, excitatory neuron subtypes; Endo, endothelia cells; Micro, microglia; Oligo, oligodendrocytes; OPC, oligodendrocyte progenitor cells. **d** Shared cell-cell communications between STARmap and Slide-seq data inferred by SpaTalk. The asterisk represents the result with at least one LRI in Slide-seq data, while the well number means the result with at least one shared LRI between STARmap and Slide-seq data. **e** Top 20 inferred LRIs that mediate the cell-cell communication from the eL6 and Pvalb neuron senders to the Oligo receivers and the significantly enriched biological processes and pathways with the inferred ligand and receptor genes using Metascape (left). The right panel is the enriched gene sets using GSEA over Oligo.

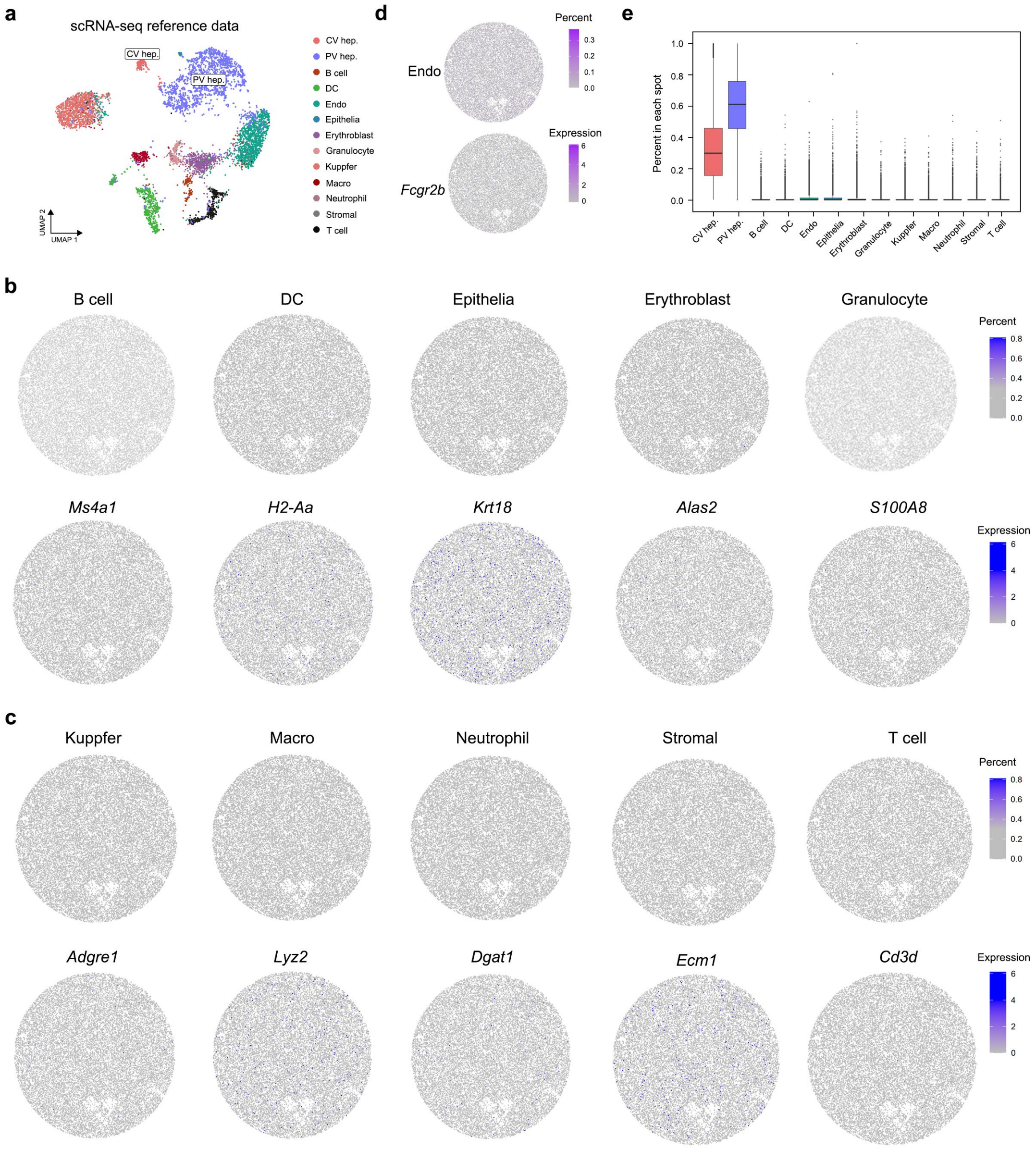


**Fig. 4 Cell-type decomposition on the mouse liver ST data of Slide-seq**. **a** Mouse liver scRNA-seq reference integrating the non-parenchymal cells from the mouse cell atlas (MCA)46 and the parenchymal hepatic cells from GSE12568847, which contains 6029 cells involving the major immune cells and the pericentral and periportal hepatocytes, etc. **b-d** Expression of known marker gene (up) and the percent (down) for Endo, B cell, DC, Epithelia, Erythroblast, granulocyte, Kupffer cell, Macro, neutrophil, stromal cell, and T cell. **e** Percent of cell types across 25595 spots of Slide-seq data.

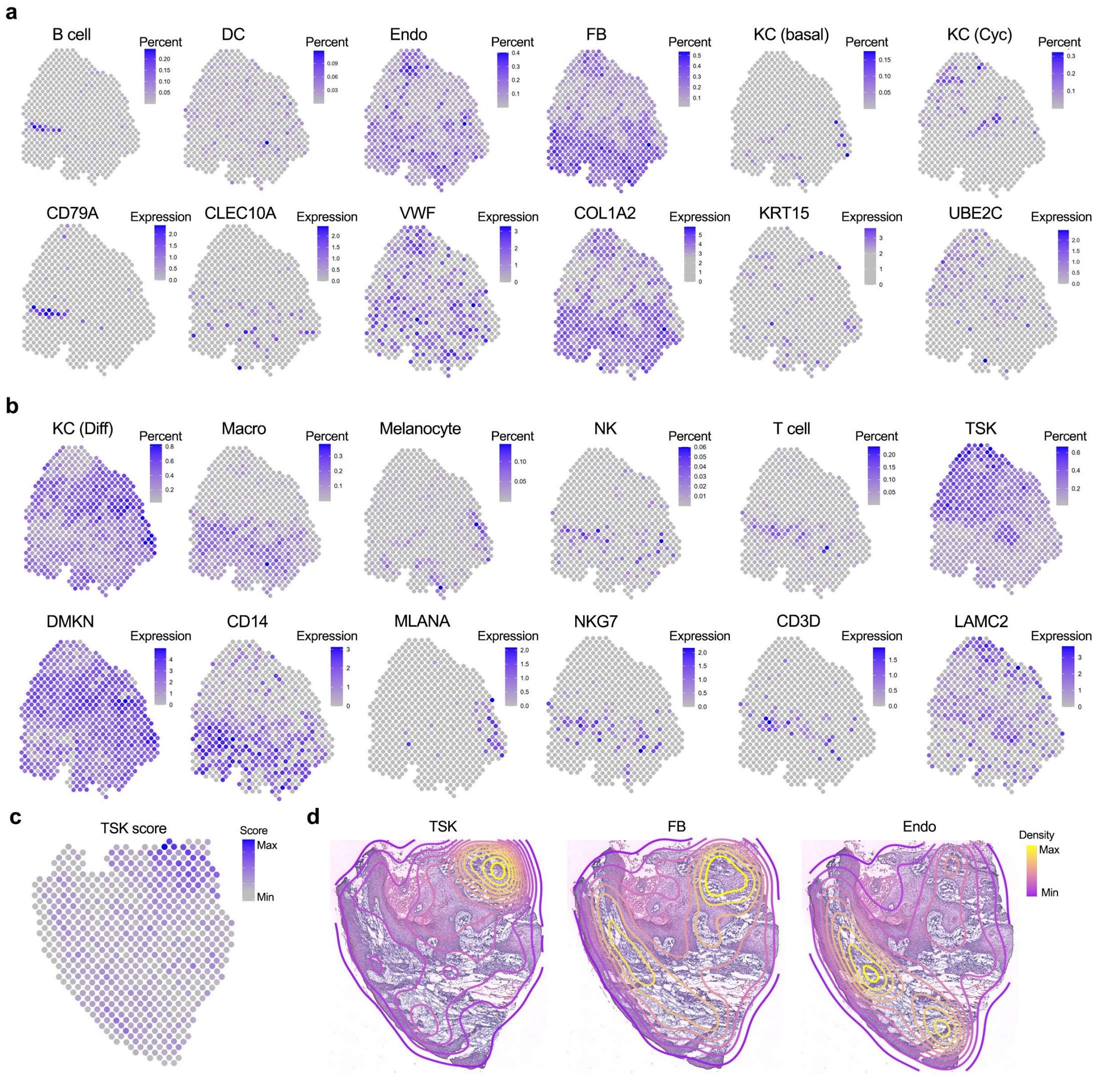


**Fig. 5 Cell-type decomposition on the human skin SCC ST data of 10X Visium**. **a-b** Expression of known marker gene (up) and the percent (down) for B cell, dendritic cell (DC), endothelial cell (Endo), fibroblast (FB), KC (basal), KC (Cycling), KC (Differentiating), macrophage (Macro), melanocyte, natural killer (NK) cell, T cell, and TSK. **c** TSK score across spatial spots using the signatures of TSK. **d** Contour plot of TSK, FB, and Endo based on the reconstructed single-cell ST atlas by SpaTalk in patient 10.

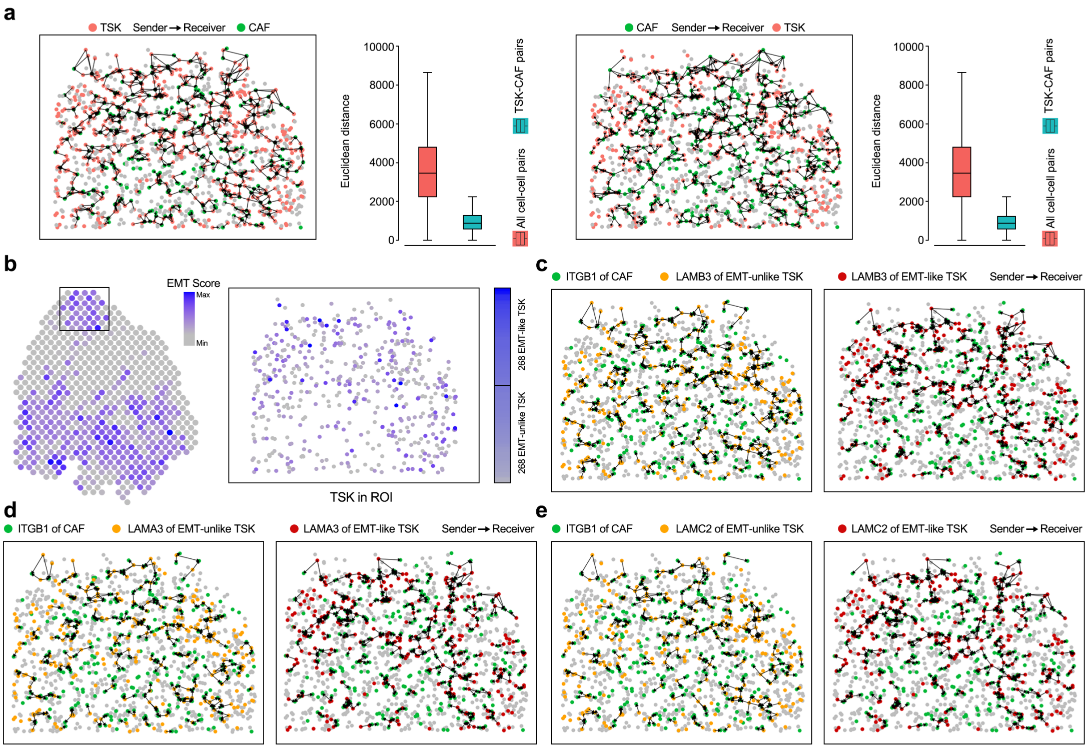


**Fig. 6 Cell-cell communications between TSK subpopulation and stromal cells in space with SpaTalk. a** Direct cell-cell communication between TSK and CAF. Neighbor cells for TSK and CAF senders were plotted, respectively, wherein the left are the cell-cell pairs of TSK senders and CAF receivers and the right are the cell-cell pairs of CAF senders and TSK receivers. **b** EMT score across spatial spots using the signatures of EMT hallmarks, wherein TSKs were divided into EMT-like and EMT-unlike TSKs according to the EMT scores. **c-e** Number of cell-cell pairs from the EMT-like and EMT-unlike TSKs to CAFs over the LRIs, namely LAMB3-ITGB1, LAMA3-ITGB1, and LAMC2-ITGB1.
